# Phenotypic clustering reveals distinct subtypes of polycystic ovary syndrome with novel genetic associations

**DOI:** 10.1101/814210

**Authors:** Matthew Dapas, Frederick T. J. Lin, Girish N. Nadkarni, Ryan Sisk, Richard S. Legro, Margrit Urbanek, M. Geoffrey Hayes, Andrea Dunaif

## Abstract

**Background:** Polycystic ovary syndrome (PCOS) is a common, complex genetic disorder affecting up to 15% of reproductive age women worldwide, depending on the diagnostic criteria applied. These diagnostic criteria are based on expert opinion and have been the subject of considerable controversy. The phenotypic variation observed in PCOS is suggestive of an underlying genetic heterogeneity, but a recent meta-analysis of European ancestry PCOS cases found that the genetic architecture of PCOS defined by different diagnostic criteria was generally similar, suggesting that the criteria do not identify biologically distinct disease subtypes. We performed this study to test the hypothesis that there are biologically relevant subtypes of PCOS.

**Methods and Findings:** Unsupervised hierarchical cluster analysis was performed on quantitative anthropometric, reproductive, and metabolic traits in a genotyped discovery cohort of 893 PCOS cases and an ungenotyped validation cohort of 263 PCOS cases. We identified two PCOS subtypes: a “reproductive” group (21-23%) characterized by higher luteinizing hormone (LH) and sex hormone binding globulin (SHBG) levels with relatively low body mass index (BMI) and insulin levels; and a “metabolic” group (37-39%), characterized by higher BMI, glucose, and insulin levels with lower SHBG and LH levels. We performed a GWAS on the genotyped cohort, limiting the cases to either the reproductive or metabolic subtypes. We identified alleles in four novel loci that were associated with the reproductive subtype at genome-wide significance (*PRDM2/KAZN1*, P=2.2×10^-10^; *IQCA1*, P=2.8×10^-9^; *BMPR1B/UNC5C*, P=9.7×10^-9^; *CDH10,* P=1.2×10^-8^) and one locus that was significantly associated with the metabolic subtype (*KCNH7/FIGN*, P=1.0×10^-8^). We have previously reported that rare variants in *DENND1A*, a gene regulating androgen biosynthesis, were associated with PCOS quantitative traits in a family-based whole genome sequencing analysis. We classified the reproductive and metabolic subtypes in this family-based PCOS cohort and found that the subtypes tended to cluster in families and that carriers of rare *DENND1A* variants were significantly more likely to have the reproductive subtype of PCOS. Limitations of our study were that only PCOS cases of European ancestry diagnosed by NIH criteria were included, the sample sizes for the subtype GWAS were small, and the GWAS findings were not replicated.

**Conclusions:** In conclusion, we have found stable reproductive and metabolic subtypes of PCOS. Further, these subtypes were associated with novel susceptibility loci. Our results suggest that these subtypes are biologically relevant since they have distinct genetic architectures. This study demonstrates how precise phenotypic delineation can be more powerful than increases in sample size for genetic association studies.

## Introduction

Understanding the genetic architecture of complex diseases is a central challenge in human genetics (1–3). Often defined according to arbitrary diagnostic criteria, complex diseases can represent the phenotypic convergence of numerous genetic etiologies (4–8). Recent studies in type 2 diabetes support the concept that there are disease subtypes with distinct genetic architecture (7, 8). Identifying and addressing genetic heterogeneity in complex diseases could increase power to detect causal variants and improve treatment efficacy (9).

Polycystic ovary syndrome (PCOS) is a highly heritable, complex genetic disorder affecting up to 15% of reproductive-age women worldwide, depending on the diagnostic criteria applied (10). It is characterized by a variable constellation of reproductive and metabolic abnormalities (**Fig 1**). It is the leading cause of anovulatory infertility and a major risk factor for type 2 diabetes (T2D) in women (11). Despite these substantial morbidities, the etiology(ies) of PCOS remains unknown (12). Accordingly, the commonly-used diagnostic criteria for PCOS, the National Institutes of Health (NIH) criteria (13) and the Rotterdam criteria (14, 15), are based on expert opinion, rather than mechanistic insights, and are designed to account for the diverse phenotypic presentations of PCOS. The NIH criteria require the presence of hyperandrogenism (HA) and chronic oligo/anovulation or ovarian dysfunction (OD) (13). The Rotterdam criteria include polycystic ovarian morphology (PCOM) and require the presence of at least two of these three key reproductive traits, resulting in three different affected phenotypes: HA and OD with or without PCOM, also known as NIH PCOS, as well as two additional non-NIH Rotterdam phenotypes, HA and PCOM, and OD and PCOM.

**Fig 1.**
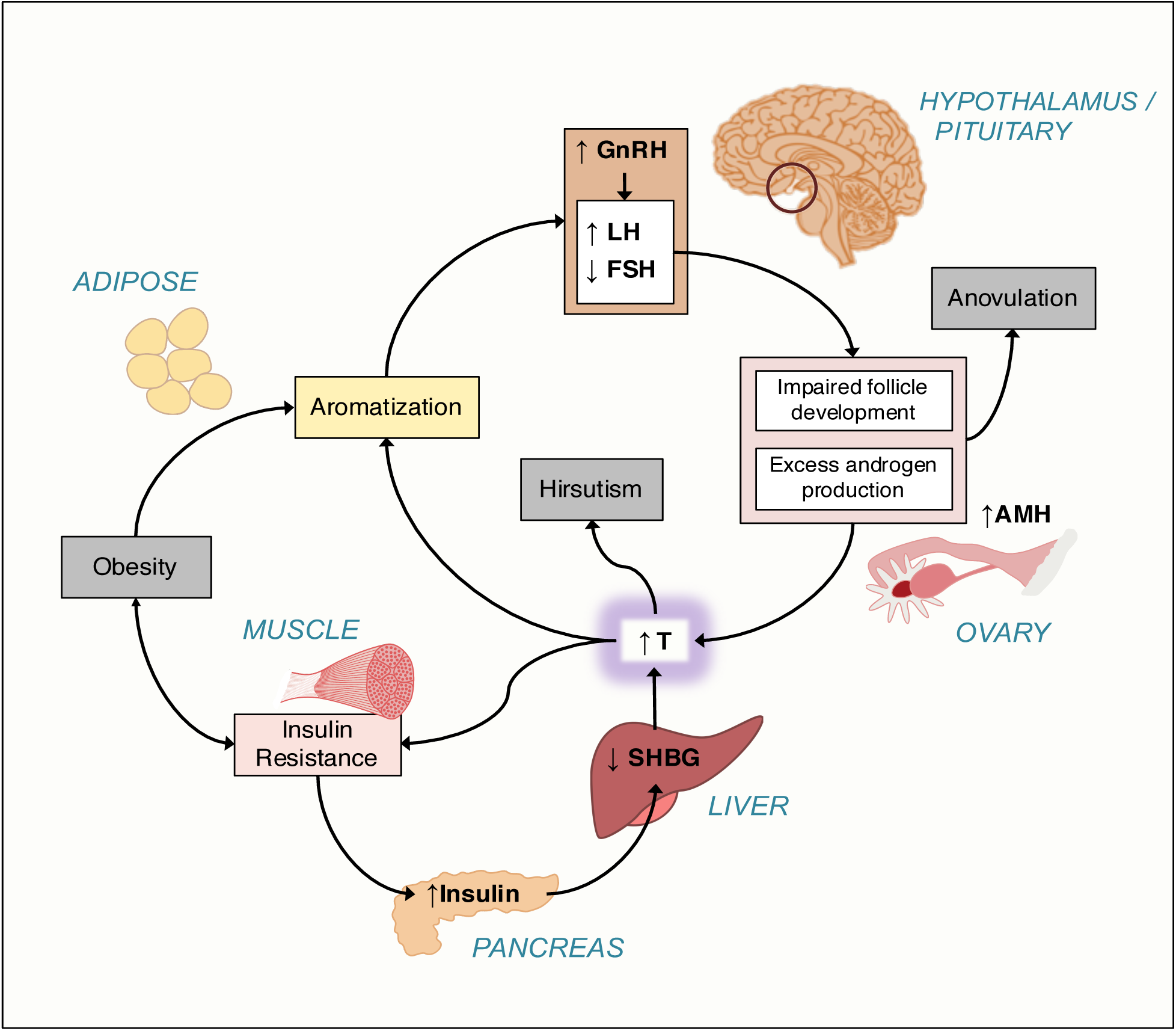
Pathophysiology of PCOS (reviewed in (10)). There is increased frequency of pulsatile gonadotropin-releasing hormone (GnRH) secretion from the arcuate nucleus of the hypothalamus that selectively increases luteinizing hormone (LH) secretion. LH stimulates ovarian theca cell testosterone (T) production. T is incompletely aromatized to estradiol by the adjacent granulosa cells because of relative follicle stimulating hormone (FSH) deficiency. There are constitutive increases in the activity of multiple steroidogenic enzymes in theca cells from women with PCOS, which contributes to increased T production. Increased adrenal androgen production may also be present in PCOS. T acts in the periphery to produce clinical signs of androgen excess, such as hirsutism, acne, and alopecia. T and androstenedione can also be aromatized extragonadally to estradiol and estrone, respectively, resulting in unopposed estrogen action on the endometrium. T feeds back on the hypothalamus to decrease the sensitivity to the normal feedback effects of estradiol and progesterone to slow GnRH pulse frequency. Anti-Müllerian hormone (AMH) levels are frequently increased in PCOS; this hormone is secreted by small, growing preantral follicles, which are increased in PCOS. Recent studies suggest AMH acting through its cognate receptor on GnRH neurons in the arcuate nucleus contributes to the pathogenesis of PCOS (16, 17). PCOS is often associated with profound insulin resistance due to a unique defect in post-binding insulin-mediated signal transduction. Insulin is a co-gonadotropin that acts in synergy with LH to amplify the reproductive abnormalities of PCOS. Insulin signaling in the hypothalamus also appears to be important for ovulation. Insulin is a major negative regulator of hepatic synthesis of sex hormone-binding globulin (SHBG), the specific transport protein for T; only T which is not bound to SHBG is biologically active.

Genomewide association studies (GWAS) have considerably advanced our understanding of the pathophysiology of PCOS. These studies have implicated gonadotropin secretion (18) and action (19, 20), androgen biosynthesis (19–21), metabolic regulation (21, 22) and ovarian aging (22) in PCOS pathogenesis. A recent meta-analysis (21) of GWAS was the first study to investigate the genetic architecture of the diagnostic criteria. Only one of 14 PCOS susceptibility loci identified was significantly more strongly associated with the NIH phenotype compared to non-NIH Rotterdam phenotypes or to self-reported PCOS. These findings suggested that the genetic architecture of the phenotypes defined by the different PCOS diagnostic criteria was generally similar. Therefore, the current diagnostic criteria do not appear to identify genetically distinct disease subtypes.

It is possible to identify physiologically relevant complex disease subtypes through cluster analysis of phenotypic traits (7,23,24). Indeed, there have been previous efforts to subtype PCOS using unsupervised cluster analysis of its hormonal and anthropometric traits (25–28). However, there has been no validation that the resulting PCOS subtypes were biologically meaningful by testing their association with genetic variants, with other independent biomarkers, or with outcomes, such as therapeutic responses. In this study, we sought to 1) identify phenotypic subtypes of PCOS using an unsupervised clustering approach on reproductive and metabolic quantitative traits from a large cohort of women with PCOS, 2) validate those subtypes in a replication cohort, and 3) test whether the subtypes thus identified were associated with distinct common genetic variants. As an additional validation, we investigated the association of the subtypes with rare genetic variants we recently identified in a family-based PCOS cohort (29).

## Methods

### Subjects

This study used biochemical and genotype data from the previously published PCOS GWAS, Hayes and Urbanek et al., 2015 (18), in which a discovery cohort (Stage 1) of 984 PCOS cases and 2,964 population controls were studied, followed by a replication cohort (Stage 2) of 1,799 PCOS cases and 1,231 phenotyped reproductively normal control women. All cases were of European ancestry and each subject provided written informed consent prior to the study (18). PCOS cases were ages 13-45 years and were diagnosed according to the NIH criteria (10) of hyperandrogenism and chronic anovulation (eight or fewer menses per year), excluding specific disorders of the adrenals, ovaries, or pituitary (30). Cases fulfilling the NIH criteria also meet the Rotterdam criteria for PCOS (10). Two additional PCOS cohorts were included in the present study who fulfilled the NIH criteria and were phenotyped according to the same methods as the genotyped GWAS cohort. An independent, ungenotyped cohort of 263 women with PCOS was used for clustering replication. A family-based whole-genome sequencing cohort of 73 women with PCOS was used for investigating subtype clustering in families and for rare variant analysis (29).

Population-based control DNA samples for the GWAS Stage 1 cohort were obtained from the NUgene biobank (31) from women of European ancestry, ages 18-97 years. Control women in the Stage 2 cohort were phenotyped reproductively normal women of European ancestry, ages 15–45 years, with regular menses and normal T levels, and who were not receiving contraceptive steroids for at least 3 months prior to study (18). T, DHEAS, SHBG, luteinizing hormone (LH), follicle-stimulating hormone (FSH), fasting glucose (Glu0), and fasting insulin (Ins0) levels were measured as previously reported (18).

### Clustering

Clustering was performed in PCOS cases on eight adjusted quantitative traits: BMI, T, DHEAS, Ins0, Glu0, SHBG, LH, and FSH. There were 893 combined cases from both stages with complete quantitative trait data available for clustering (**Table S1**). Quantitative trait values were first log_e_-normalized and adjusted for age and assay method, which varied according to the different study sites where samples were collected (18), using a linear regression. An inverse normal transformation was then applied for each trait to ensure equal scaling. The normalized trait residuals were clustered using unsupervised, agglomerative, hierarchical clustering according to a generalization of Ward’s minimum variance method (32, 33) on Manhattan distances between trait values. Differences in adjusted, normalized trait values between subtypes were assessed using Kruskal-Wallis and pairwise Wilcoxon rank-sum tests corrected for multiple testing (Bonferroni). Cluster stability was assessed by computing the mean Jaccard coefficient from a repeated nonparametric bootstrap resampling (n=1000) of the dissimilarity matrix (34). Jaccard coefficients below 0.5 indicate that a cluster does not capture any discernable pattern within the data, while a mean coefficient above 0.6 indicates that the cluster reflects a real pattern within the data (34). Cluster reproducibility was further assessed by repeating the clustering procedure in an independent cohort of 263 PCOS cases.

### Association Testing

Stage 1 samples were genotyped using the Illumina OmniExpress (HumanOmniExpress-12v1_C) single nucleotide polymorphism (SNP) array. Stage 2 samples were genotyped using the Metabochip (35) with added custom variant content based on ancillary studies and the discovery results (18). The Stage 2 association replication in this study was therefore limited; many of the loci from Stage 1 were therefore not characterized in Stage 2. Low quality genotypic data were removed as described previously (18). SNPs were filtered according to minor allele frequency (MAF ≥0.01), Hardy-Weinberg equilibrium (p ≥1×10^-6^), call rate (≥0.99), minor allele count (MAC>5), Mendelian concordance, and duplicate sample concordance. Only autosomal SNPs were considered. Ancestry was evaluated using a principal component analysis (PCA) (36) on 76,602 LD-pruned SNPs (18). Samples with values >3 standard deviations from the median for either of the first two principal components (PCs) were excluded (34 in discovery; 37 in replication). Genotype data was phased using ShapeIT (v2.r790) (37) and then imputed to the 1000 Genomes reference panel (Phase3 v5) (38) using Minimac3 via the Michigan Imputation Server (39). Imputed SNPs with an allelic r^2^ below 0.8 were removed from analysis.

Association testing was performed separately for Stage 1 and Stage 2 cohorts. Of the 893 combined cases from both stages included in the clustering analysis, 555 were from the Stage 1 cohort and 338 were from the Stage 2 cohort. 2,964 normal controls were used in Stage 1, and 1,134 were used in Stage 2. Logistic regressions were performed using SNPTEST (40) for case-control status under an additive genetic model, adjusting for BMI and first three PCs of ancestry. P-values are reported as P_1_ and P_2_ for Stage 1 and Stage 2, respectively. Cases were limited to specific subtypes selected from clustering results. The betas and standard errors were combined across Stage 1 and Stage 2 cohorts for each subtype under a fixed meta-analysis model weighting each strata by sample size (41). Association test outputs were aligned to the same reference alleles and weighted z-scores were computed for each SNP. The square root of the cohort-specific sample size was used as the proportional weight. Meta-analysis P-values (P_meta_) were adjusted for genomic inflation. Associations with P-values < 1.67×10^-8^ were considered statistically significant, based on the standard P<5×10^−8^ used in conventional GWAS adjusted for the three independent association tests performed.

### Chromatin interactions

Neighboring chromatin interactions were investigated in intergenic loci using high-throughput chromatin conformation capture (Hi-C) data from the 3DIV database (42). Topologically associating domains (TADs) were identified using TopDom (43) with a window size of 20.

### Identifying subtypes in PCOS families

Quantitative trait data from the affected women (n=73) in the family-sequencing cohort (29) were adjusted and normalized as described above. Subtype classifiers were modeled on the adjusted trait values and cluster assignments from the genotyped cohort. Several classification methods were compared using 10-fold cross-validation, including support vector machine, random forest (RF), Gaussian mixed-model, and quadratic discriminant analysis (44). The classifier with the lowest error rate was then applied to the affected women in the family-sequencing cohort to identify subtypes of PCOS in the family data. Some of the probands from the family-based cohort were included in our previous GWAS (18). Therefore, there was some sample overlap between the training and test cohorts: of the 893 genotyped women used to identify the original subtype clusters, 47 were also probands in the family-based cohort. Differences between subtypes in the proportion of women with *DENND1A* rare variants were tested using the chi-square test of independence.

## Results

### PCOS subtypes

The clustering revealed two distinct phenotypic subtypes: 1) a group (23%) characterized by higher LH and SHBG levels with relatively low BMI and Ins0 levels, which we designated “reproductive”, and 2) a group (37%) characterized by higher BMI and Glu0 and Ins0 levels with relatively low SHBG and LH levels, which we designated “metabolic” (**Fig 2**). The key traits distinguishing the reproductive and metabolic subtypes were BMI, insulin, SHBG, glucose, LH, and FSH, in order of importance according to relative pairwise Wilcoxon rank-sum test statistics (**Fig 3**). The remaining cases (40%) demonstrated no distinguishable pattern regarding their relative phenotypic trait distributions and were designated “indeterminate”. The reproductive and metabolic subtypes clustered along opposite ends of the SHBG vs. Ins0/BMI axis, which was highly correlated with the first PC of the adjusted quantitative traits (**Fig 4**). The reproductive subtype was the most stable cluster, with a mean bootstrapped Jaccard coefficient (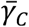) of 0.61, followed by the metabolic subtype with 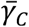=0.55. The indeterminate group did not appear to capture any discernable pattern within the data (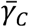=0.41) and was both overlapping and intermediate between the reproductive and metabolic subtypes on the SHBG vs. Ins0/BMI axis.

**Fig 2.**
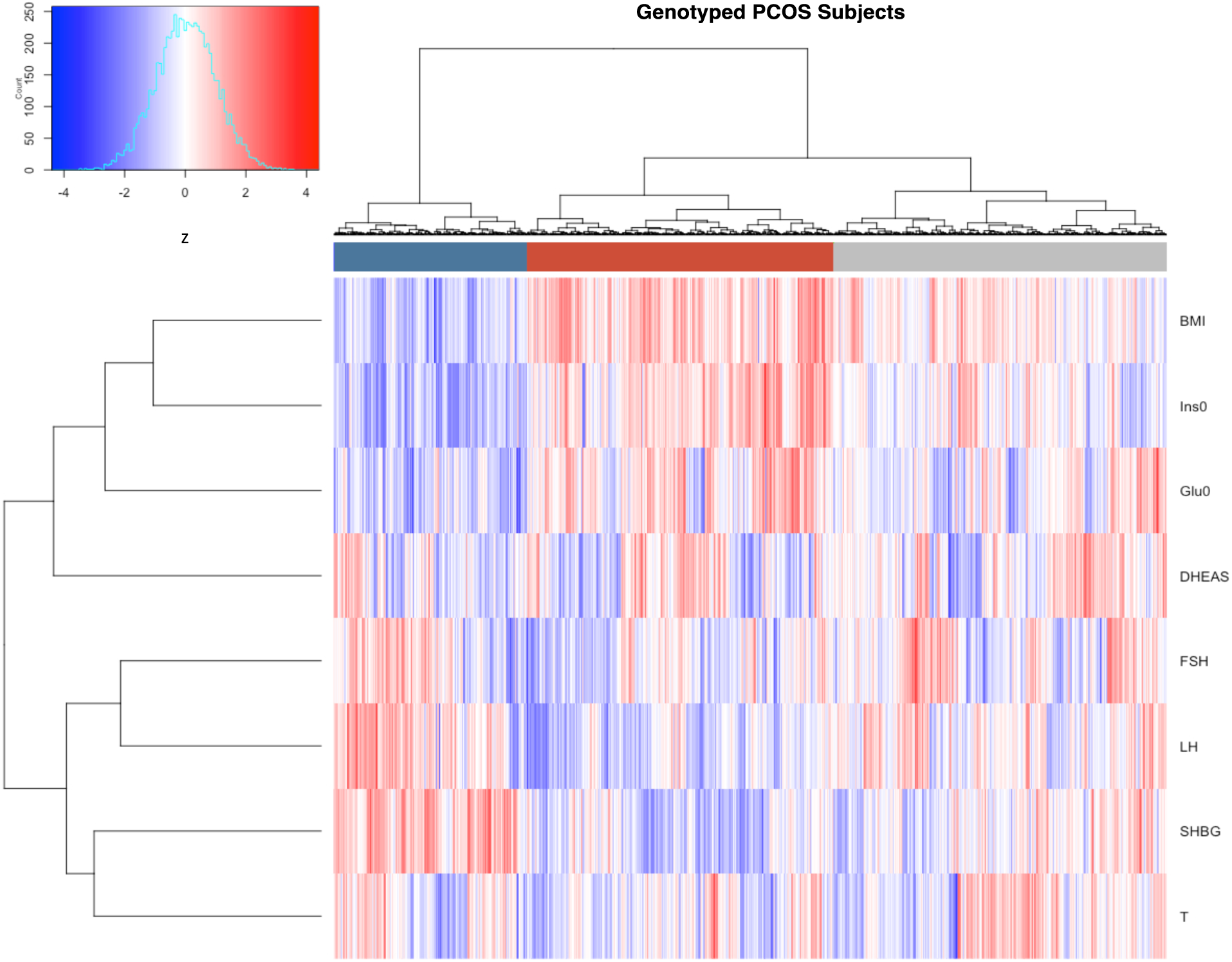
Hierarchical clustering of genotyped PCOS cases. Hierarchical clustering of 893 genotyped PCOS cases according to adjusted quantitative traits revealed two distinct phenotypic subtypes: a “reproductive” cluster, and a “metabolic” cluster; the remaining cases were designated as “indeterminate”. The reproductive, metabolic, and indeterminate clusters are shown in the color bar as dark blue, dark red, and grey, respectively. Heatmap colors correspond to trait z-scores, as shown in the frequency histogram where red indicates high values and blue indicates low values for each trait. BMI, body mass index; SHBG, sex hormone binding globulin; DHEAS, dehydroepiandrosterone sulfate; Glu0, fasting glucose; Ins0, fasting insulin; LH, luteinizing hormone; FSH, follicle-stimulating hormone.

**Fig 3.**
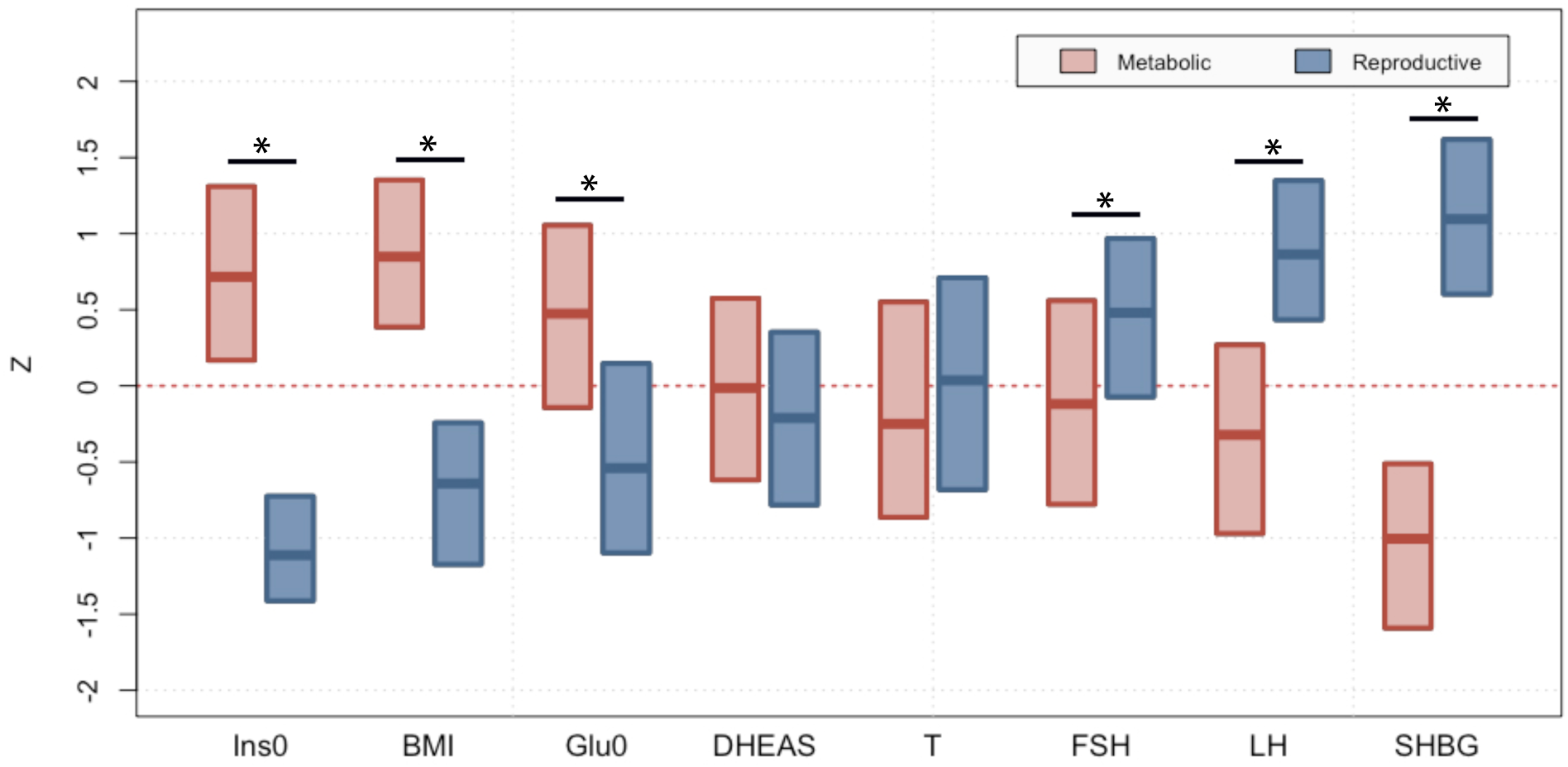
Phenotypic trait distributions in reproductive and metabolic subtypes. Median and interquartile ranges are shown for normalized, adjusted quantitative trait distributions of genotyped PCOS cases with reproductive or metabolic subtype. The figure illustrates the traits for which the subtypes differ significantly with an asterisk (*Bonferroni adjusted Wilcoxon, P_adj_<0.05): Ins0, BMI, Glu0, FSH, LH and SHBG. BMI, body mass index; SHBG, sex hormone binding globulin; DHEAS, dehydroepiandrosterone sulfate; Glu0, fasting glucose; Ins0, fasting insulin; LH, luteinizing hormone; FSH, follicle-stimulating hormone.

**Fig 4.**
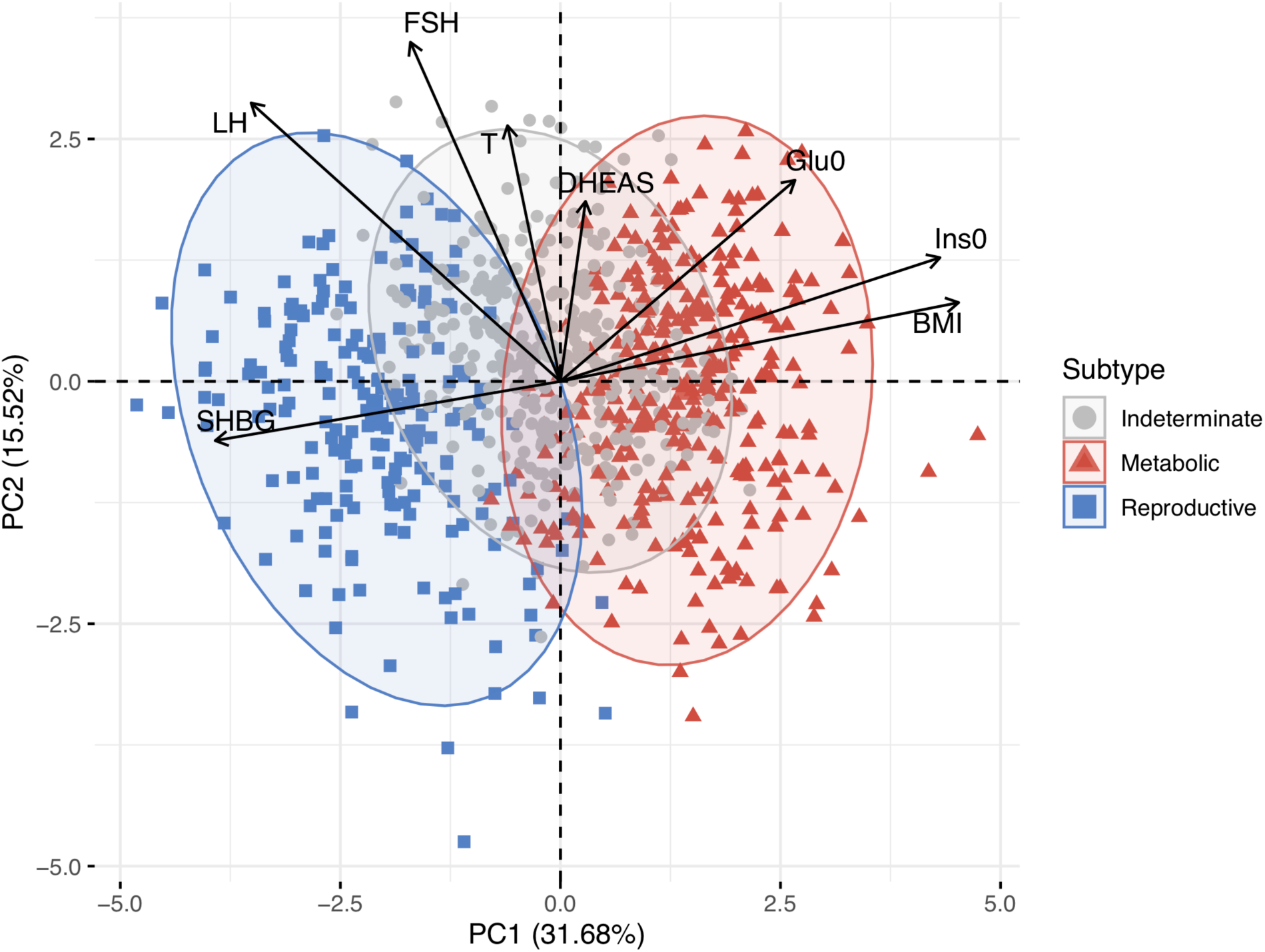
PCA plot of quantitative traits for genotyped PCOS cases. Genotyped PCOS cases are plotted on the first two principal components of the quantitative trait data and colored according to their identified subtype. Subtype clusters are shown as 95% concentration ellipses, assuming bivariate normal distributions. The relative magnitude and direction of trait correlations with the principal components are shown with black arrows.

The clustering procedure was then repeated in an independent, non-genotyped cohort of 263 NIH PCOS cases diagnosed according to the same criteria as the genotyped cohort. The clustering yielded similar results, with a comparable distribution of reproductive (26%, 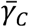=0.57), metabolic (39%, 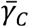=0.46), and indeterminate clusters (35%, 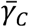=0.40) (**Fig 5**).

**Fig 5.**
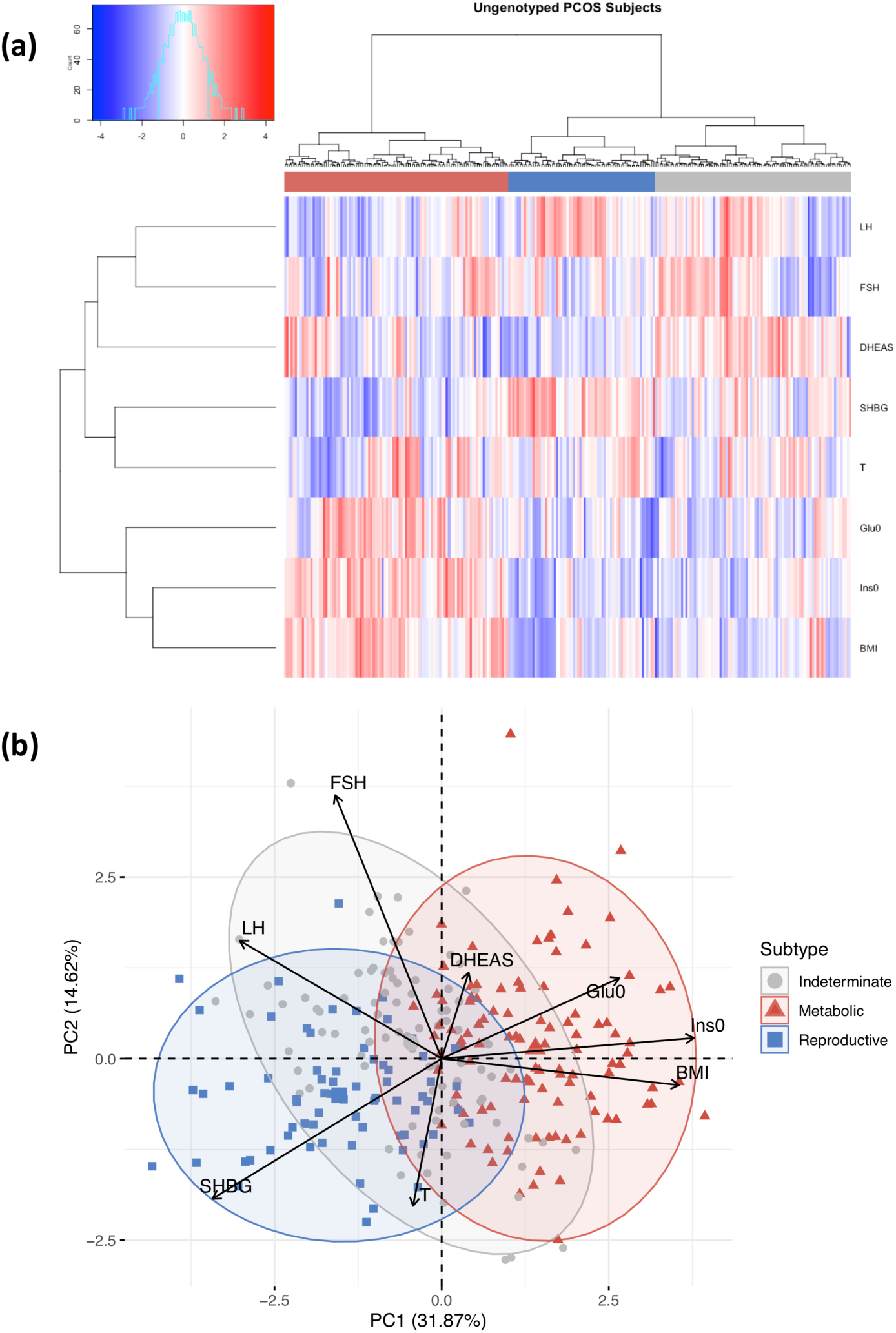
Clustering of ungenotyped PCOS cases. (a) Hierarchical clustering of 263 ungenotyped PCOS cases according to adjusted quantitative traits replicate reproductive (blue), metabolic (red), and unclassified (grey) clusters. Heatmap colors correspond to trait z-scores. (b) PCA plot of ungenotyped PCOS cases replicate results from genotyped cases. (a) Hierarchical clustering of 263 ungenotyped PCOS cases according to adjusted quantitative traits replicate reproductive (blue), metabolic (red), and unclassified (grey) clusters. Heatmap colors correspond to trait z-scores. (b) PCA plot of ungenotyped PCOS cases replicate results from genotyped cases.

### Subtype genetic associations

Genome-wide association testing identified alleles in four novel loci that were associated with the reproductive PCOS subtype at genome-wide significance (chr1 p36.21 *PRDM2/KAZN*, P=2.23×10^-10^; chr2 q37.3 *IQCA1*, P=2.76×10^-9^; chr4 q22.3 *BMPR1B/UNC5C*, P=9.71×10^-9^; chr5 p14.2-p14.1 *CDH10,* P=1.17×10^-8^) and one novel locus that was significantly associated with the metabolic subtype (chr2 q24.2-q24.3 *KCNH7/FIGN*, P=1.03×10^-8^). Association testing on the indeterminate cases replicated the 11p14.1 *FSHB* locus from our original GWAS (18) (**Table 1; Figs 6 and 7**).

**Table 1.** Genome-wide significant associations with PCOS subtypes.

| Chr | Mb | Variant | Gene(s) | EA | Stage 1 (Discovery) | | | | | | Stage 2 (Replication) | | | | | | $P_{meta}$ |
| --- | --- | --- | --- | --- | --- | --- | --- | --- | --- | --- | --- | --- | --- | --- | --- | --- | --- |
| | | | | | EAF | $\beta$ | OR | 95% CI | P | Imp $r^2$ | EAF | $\beta$ | OR | 95% CI | P | Imp $r^2$ | |
| 1 | 14.7 | rs78025940 | *PRDM2/*<br>*KAZN* | A | 0.02 | 3.02 | 4.75 | 2.82-7.98 | $2.16 \times 10^{-10}$ | 0.91 | - | - | - | - | - | - | $2.23 \times 10^{-10}$ |
| 2 | 237.4 | rs76182733 | *IQCA1* | G | 0.01 | 3.79 | 5.68 | 3.00-10.78 | $2.67 \times 10^{-9}$ | 0.84 | - | - | - | - | - | - | $2.76 \times 10^{-9}$ |
| 4 | 96.1 | rs17023134 | *BMPR1B/*<br>*UNC5C* | G | 0.05 | 1.62 | 3.02 | 2.06-4.42 | $1.40 \times 10^{-8}$ | 0.87 | 0.06 | 0.61 | 1.71 | 0.98-2.99 | $7.81 \times 10^{-2}$ | 0.83 | $9.71 \times 10^{-9}$ |
| 5 | 24.7 | rs7735176 | *CDH10* | A | 0.01 | 3.80 | 5.09 | 2.62-9.86 | $1.14 \times 10^{-8}$ | 0.93 | - | - | - | - | - | - | $1.17 \times 10^{-8}$ |
| 2 | 164.2 | rs55762028 | *KCNH7/*<br>*FIGN* | C | 0.01 | 5.05 | 1.86 | 0.92-3.75 | $9.17 \times 10^{-9}$ | 0.96 | - | - | - | - | - | - | $1.03 \times 10^{-8}$ |
| 11 | 30.3 | rs10835638 | *FSHB* | T | 0.16 | 0.78 | 1.81 | 1.44-2.27 | $3.13 \times 10^{-8}$ | 0.98 | 0.17 | 0.77 | 2.01 | 1.49-2.70 | $2.67 \times 10^{-5}$ | 0.97 | $4.94 \times 10^{-12}$ |

Variant information and association statistics are shown for the most strongly associated SNP in each significant locus. Reproductive subtype loci are highlighted in blue, metabolic loci in red, indeterminate loci in grey. EA = effect allele; EAF: effect allele frequency in cases and controls combined; $\beta$ = effect size from association regression; OR = odds ratio; CI = confidence interval; Imp $r^2$ = imputation $r^2$ for imputed SNPs; P = stage-specific significance as assessed by logistic regression; $P_{meta}$ = significance as assessed by sample-size weighted two-strata meta-analysis, adjusted for genomic inflation factor. Cases and controls by stage: Stage 1 = 201 metabolic, 123 lean, 231 indeterminate, 2964 controls; Stage 2 = 128 metabolic, 84 lean, 126 indeterminate, 1134 controls. NOTE: Not all SNPs were genotyped or imputed in both stages.

**Fig 6.**
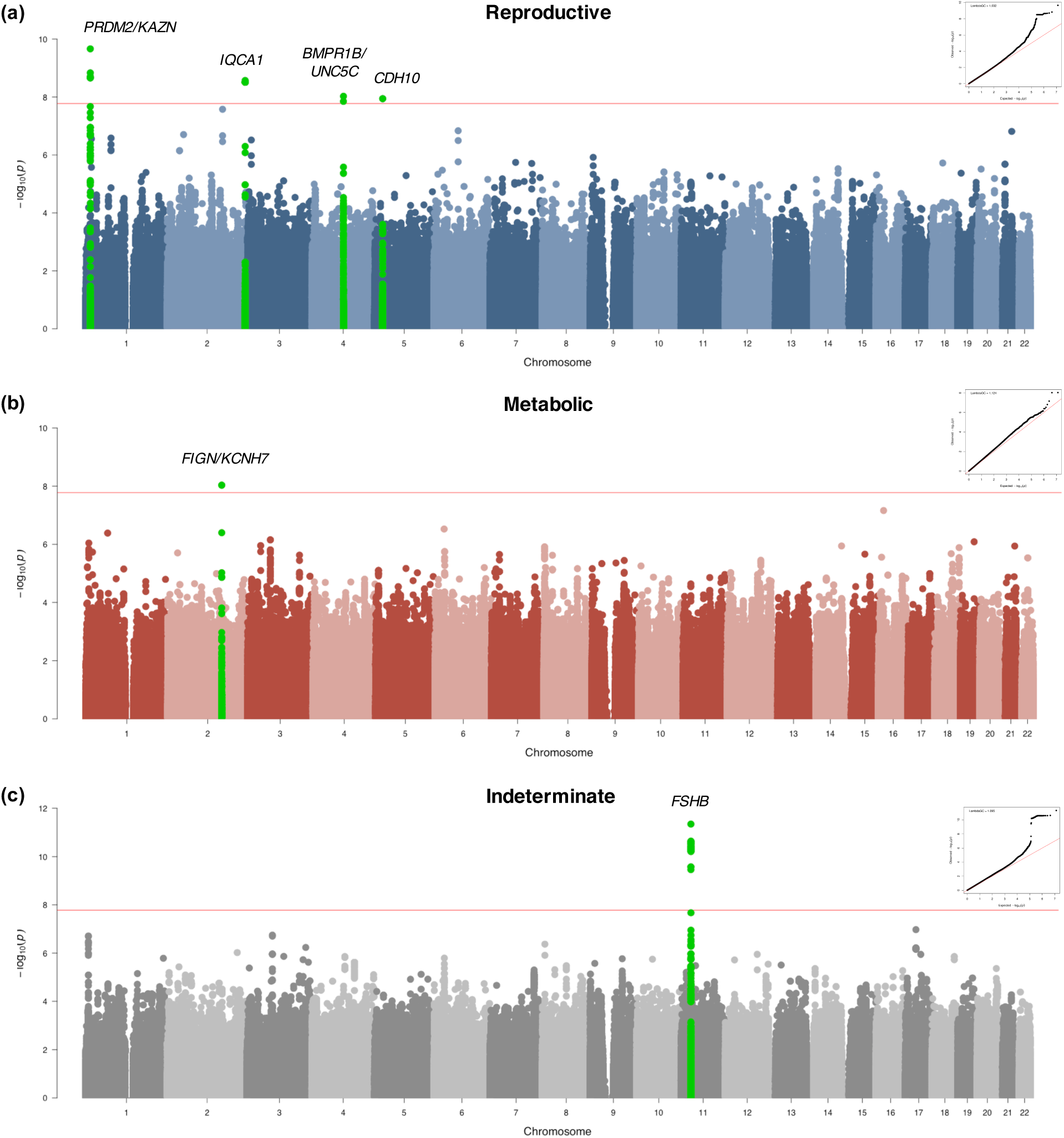
Genome-wide association results. Manhattan plots for (a) reproductive and (b) metabolic PCOS subtypes. The red horizontal line indicates genome-wide significance (p ≤ 2.5×10^-8^). Genome-wide significant loci are colored in green and labeled according to nearby gene(s). QQ plots with genomic inflation factor, λ_GC_, are shown adjacent to corresponding plots.

**Fig 7.**
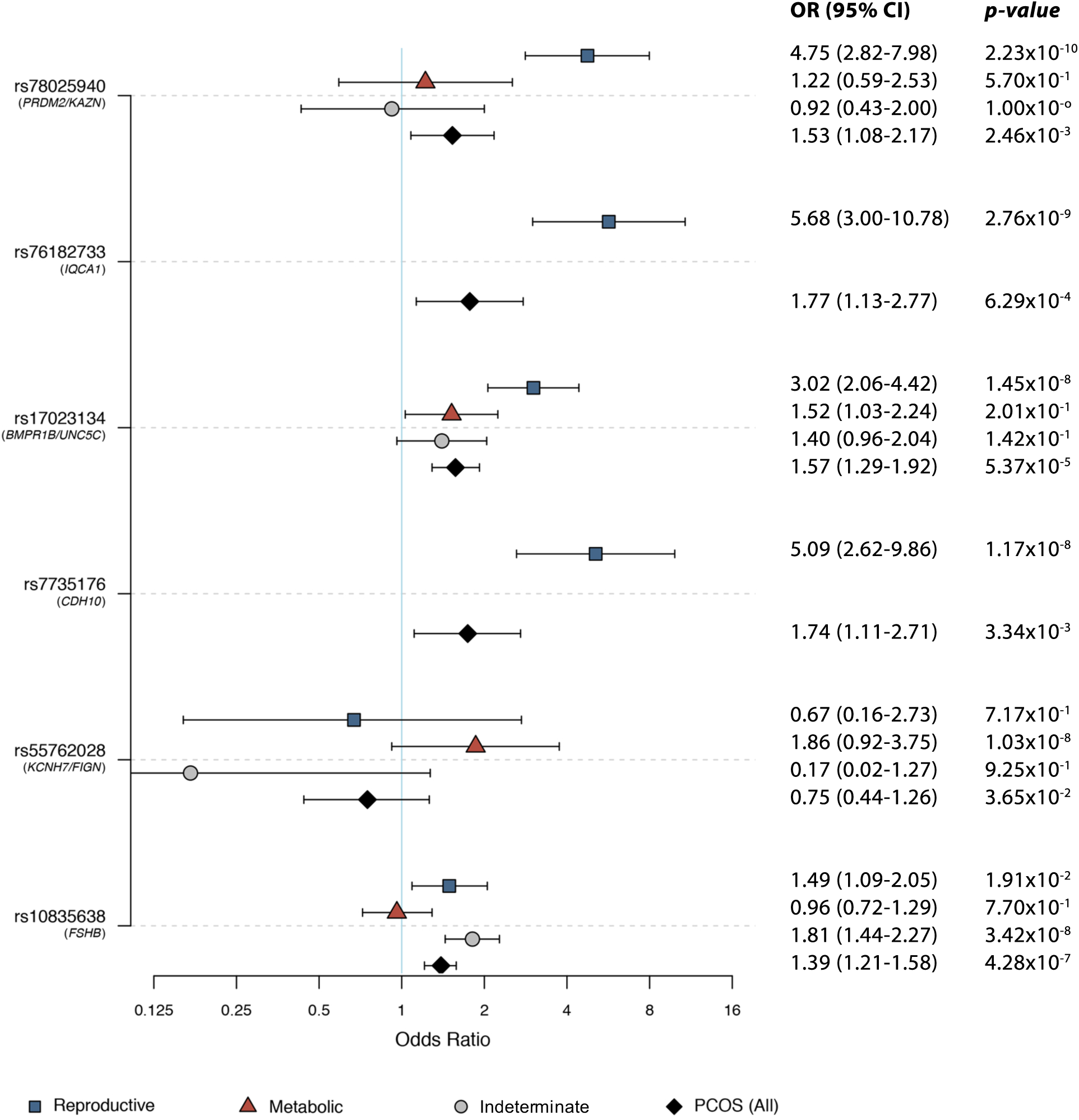
Risk allele odds ratios in PCOS and PCOS subtypes. Odds ratios with 95% confidence intervals and association P-values from the Stage 1 discovery cohort are shown for each subtype-specific novel risk allele identified in this study relative to the corresponding values for the other subtypes and for PCOS disease status in general (includes all subtypes). Some SNPs were not characterized in certain subtypes due to low allele counts or low imputation confidence.

The strongest association signal with the reproductive subtype appeared in an intergenic region of 1p36.21 579kb downstream of the *PRMD2* gene and 194kb upstream from the *KAZN* gene (**Fig 8a**). The lead SNP in the locus (rs78025940; OR=4.75, 2.82-7.98 95%CI, P_1_=2.16×10^−10^, P_meta_=2.23×10^-10^) was imputed (r^2^ = 0.91) in Stage 1 only. The SNP was not genotyped in Stage 2. The lead genotyped SNP in the locus (rs16850259) was also associated with the reproductive subtype with genome-wide significance (P_meta_=2.14×10^-9^) and was genotyped only in Stage 1 (OR=5.57, 3.24-9.56 95% CI, P_1_=2.08×10^−9^). In ovarian tissue, the SNPs appear to be centrally located within a large 2Mb TAD stretching from the *FHAD1* gene to upstream of the *PDPN* gene (**Fig 9**).

**Fig 8.**
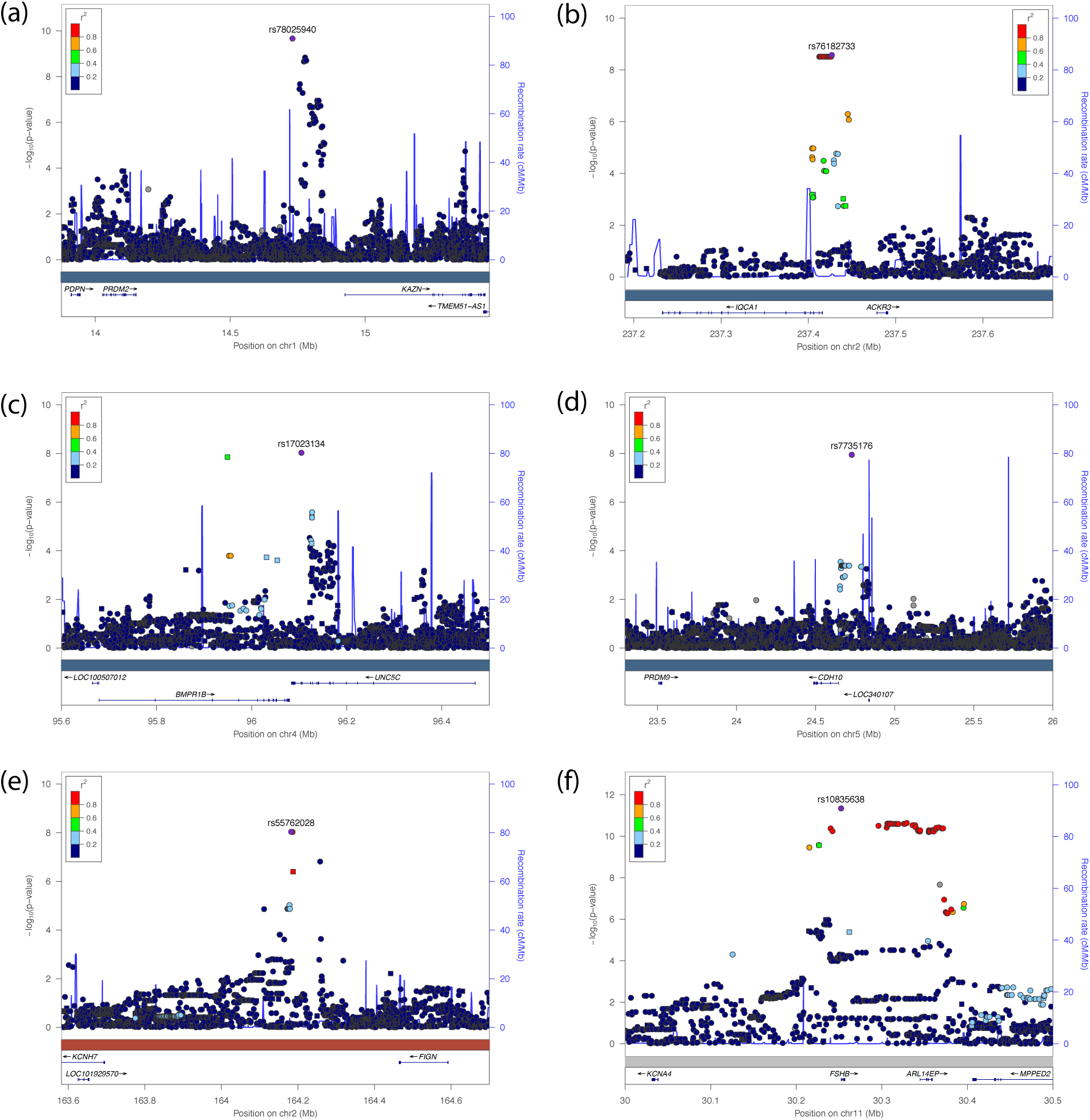
Regional association plots of genome-wide significant loci. Regional plots of association (left y-axis) and recombination rates (right y-axis) for the chromosomes (a) 1p36.21, (b) 2q37.3, (c) 4q22.3, (d) 5p14.2-p14.1, (e) 2p24.2-q24.3, and (f) 1p14.1 loci after imputation. The lead SNP in each locus is labeled and marked in purple. All other SNPs are color coded according to the strength of LD with the top SNP (as measured by r^2^ in the European 1000 Genomes data). Imputed SNPs are plotted as circles and genotyped SNPs as squares.

**Fig 9.**
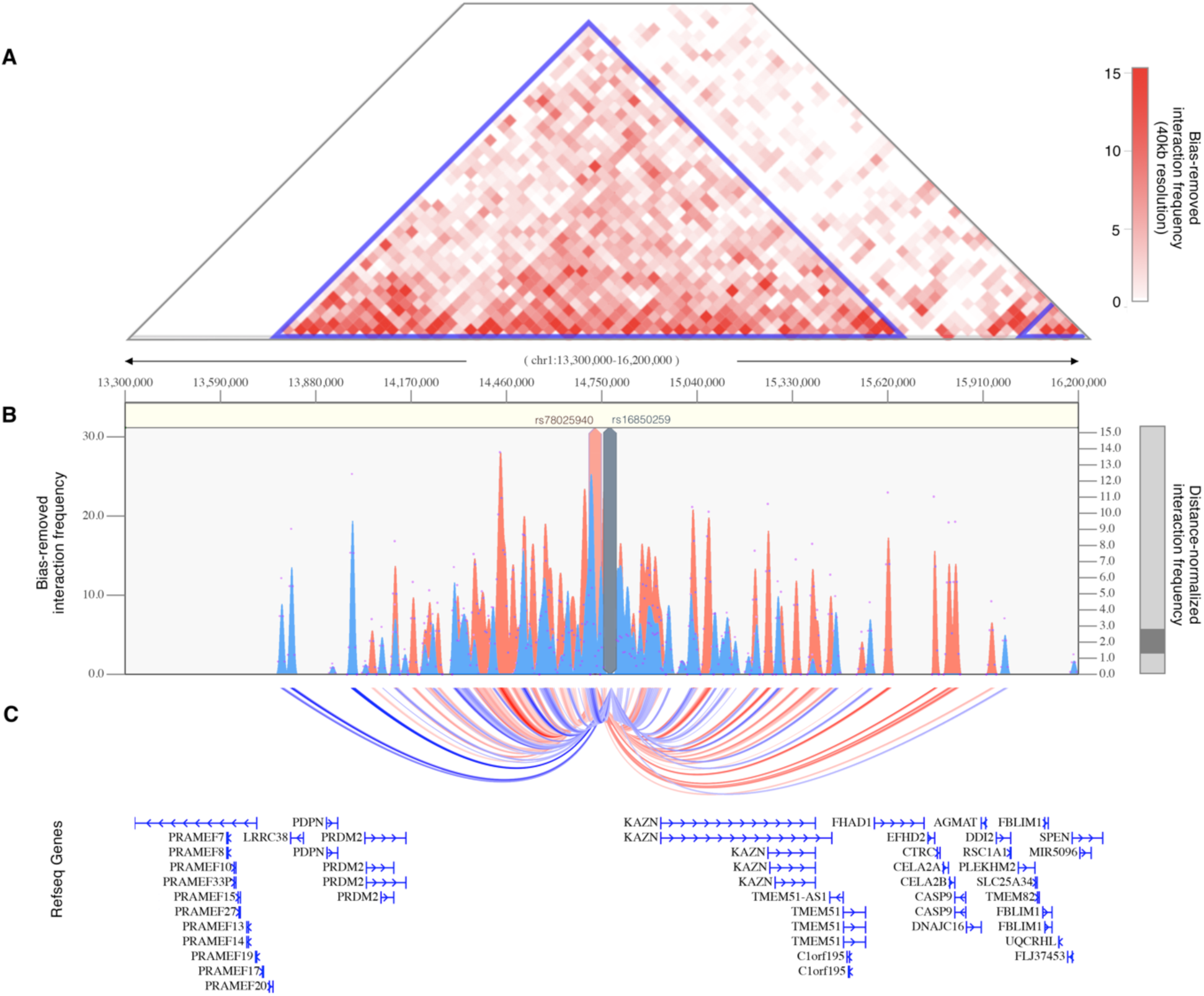
Chromatin interaction map of *PRDM2*/*KAZN* locus. (A) Shown is the interaction frequency heatmap from chr1:13,300,000-16,200,000 in ovarian tissue. The color of the heatmap indicates the level of normalized interaction frequencies with blue triangles indicating topological association domains. (B) One-to-all interaction plots are shown for the lead SNP (rs78025940; shown in red) and lead genotyped SNP (rs16850259; shown in blue) as bait. Y-axes on the left and the right measure bias-removed interaction frequency (red and blue bar graphs) and distance-normalized interaction frequency (magenta dots), respectively. (C) The arc-representation of significant interactions for distance-normalized interaction frequencies ≥ 2 is displayed relative to the RefSeq-annotated genes in the locus.

The 2q37.3 locus spanned a 50kb region of strong LD overlapping the 5’ end and promoter region of the *IQCA1* gene (**Fig 8b**). The SNP rs76182733 had the strongest association in this locus (P_meta_=2.76×10^-9^) with the reproductive subtype. The signal was genotyped only in Stage 1 (OR=5.68, 3.00-10.78 95%CI, P_1_=2.69×10^−9^) and was imputed with an imputation r^2^ value of 0.84.

The 4q22.3 locus spanned a 500kb region of LD including the 3’ ends of both the *BMPR1B* and *UNC5C* genes (**Fig 8c**). The most strongly associated SNP (rs17023134; P_meta_=9.71×10^-9^) in the locus was within an intronic region of *UNC5C*, and was associated with the reproductive subtype in the Stage 1 discovery cohort with genome-wide significance (OR=3.02, 2.06-4.42 95%CI, P_1_=1.40×10^−8^), but was not significantly associated in the Stage 2 replication analysis (OR=1.71, 0.98-2.99 95%CI, P_2_=7.8×10^−2^). The SNP was imputed with an r^2^ of 0.87 and 0.83 in the Stage 1 and Stage 2 analyses, respectively. The most strongly associated genotyped SNP in the locus (rs10516957) was also genome-wide significant (P_meta_=1.46×10^-8^) and was located in an intronic region of *BMPR1B*. The genotyped SNP was nominally associated with the reproductive subtype in both the Stage 1 (OR=2.42, 1.66-3.52 95%CI, P_1_=6.72×10^−6^) and Stage 2 (OR=2.40, 1.51-3.82 95%CI, P_2_=4.7×10^−4^) analyses with nearly identical effect sizes.

In the 5p14.2-p14.1 locus, 83kb upstream of the *CDH10* gene (**Fig 8d**), two adjacent SNPs (rs7735176, rs16893866) in perfect LD were equally associated with the reproductive subtype with genome-wide significance (P_meta_=1.17×10^-8^). The SNPs were imputed in Stage 1 (OR=5.09, 2.62-9.86 95%CI, P_1_=1.14×10^−8^) with an imputation r^2^ of 0.93.

The single locus containing genome-wide significant associations with the metabolic subtype was in an intergenic region of 2q24.2-q24.3 roughly 200kb downstream from *FIGN* and 500kb upstream from *KCNH7* (**Fig 8e**). The lead SNP, rs55762028, was imputed in Stage 1 only (OR=1.86, 0.92-3.75 95%CI, P_1_=9.17×10^−9^, P_meta_=1.03×10^-8^). In pancreatic tissue, the lead SNPs appear to be located terminally within a 1.3Mb TAD encompassing the *FIGN* gene and reaching upstream to the *GRB14* gene (**Fig 10**).

**Fig 10.**
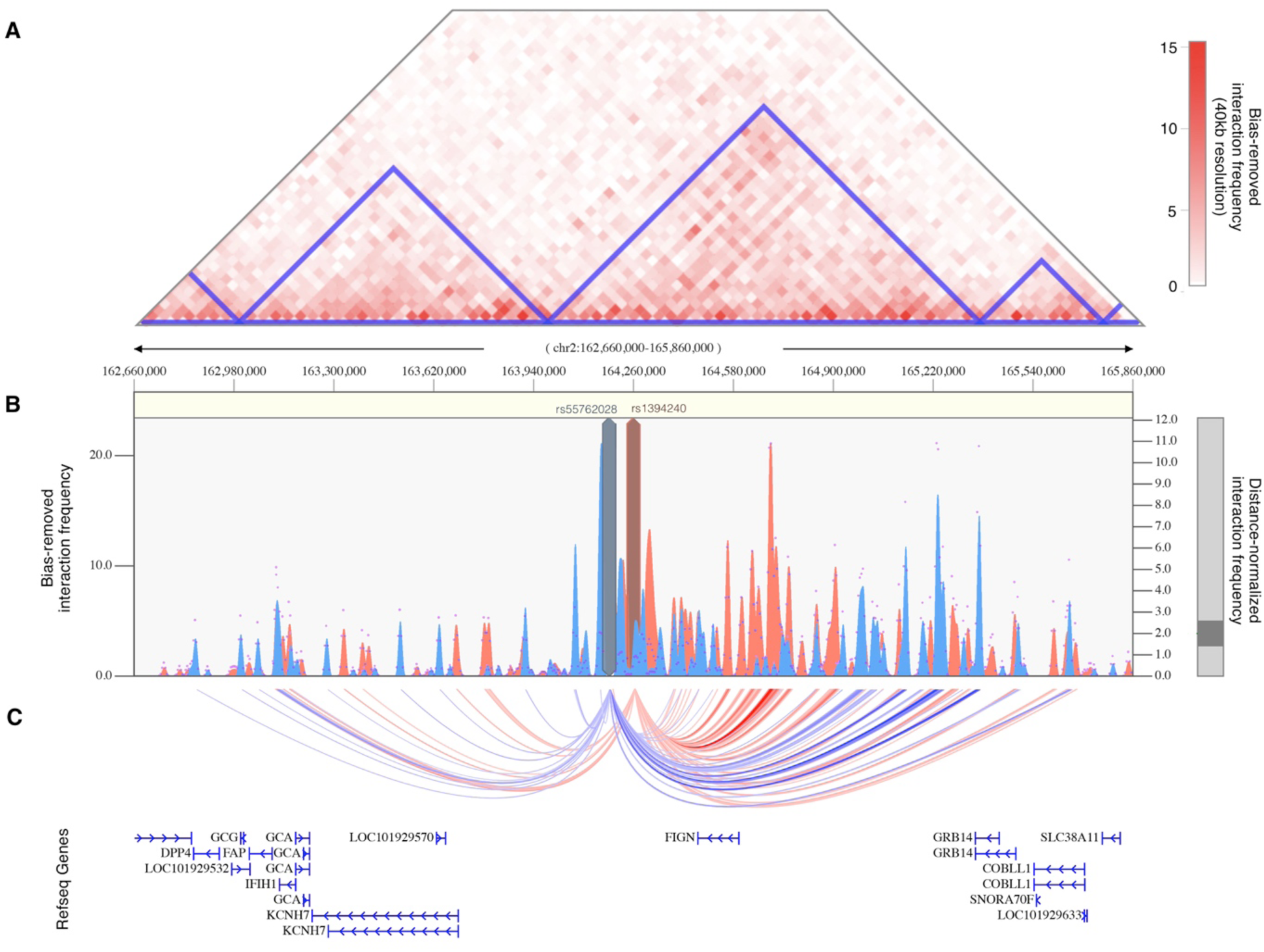
Chromatin interaction map of *KCHN7*/*FIGN* locus. (A) Shown is the interaction frequency heatmap from chr2:162,660,000 to 165,860,000 in pancreatic tissue. The color of the heatmap indicates the level of normalized interaction frequencies with blue triangles indicating topological association domains. (B) One-to-all interaction plots are shown for the lead SNP (rs13401392; shown in blue) and 2^nd^-leading SNP (rs1394240; shown in red) as bait. Y-axes on the left and the right measure bias-removed interaction frequency (blue and red bar graphs) and distance-normalized interaction frequency (magenta dots), respectively. (C) The arc-representation of significant interactions for distance-normalized interaction frequencies ≥ 2 is displayed relative to the RefSeq-annotated genes in the locus.

Association testing on the indeterminate cases replicated the genome-wide significant association in the 11p14.1 *FSHB* locus (**Fig 8f**) identified in our original GWAS (14). The lead SNP (rs10835638; P_meta_=4.94×10^-12^) and lead genotyped SNP (rs10835646; P_meta_=2.75×10^-11^) in this locus differed from the index SNPs identified in our original GWAS (rs11031006) and in the PCOS meta-analysis (rs11031005), but both of the previously identified index SNPs were also associated with the indeterminate subgroup with genome-wide significance in this study (rs11031006: P_meta_=2.96×10^-10^; rs11031005: P_meta_=2.91×10^-10^) and are in LD with the lead SNP rs10835638 (r^2^ = 0.59) (38). The other significant signals from our original GWAS (18) were not reproduced in any of the subtype association tests performed in this study (**Table 2**).

**Table 2.** Previous GWAS association signals in PCOS subtypes.

| Variant | Locus | PCOS | Reproductive | Metabolic | Indeterminate |
| --- | --- | --- | --- | --- | --- |
| rs804279 | *GATA4/NEIL2* | $P = 8.0 \times 10^{-10}$ | $P = 2.4 \times 10^{-3}$ | $P = 9.9 \times 10^{-2}$ | $P = 3.1 \times 10^{-3}$ |
| rs10993397 | *C9orf3* | $P = 4.6 \times 10^{-13}$ | $P = 2.3 \times 10^{-4}$ | $P = 6.9 \times 10^{-5}$ | $P = 1.1 \times 10^{-5}$ |
| rs11031006 | *FSHB* | $P = 1.9 \times 10^{-8}$ | $P = 8.8 \times 10^{-6}$ | $P = 6.6 \times 10^{-1}$ | $P = 3.0 \times 10^{-10}$ |

Subtype-specific association statistics are shown for each of the SNPs that were significantly associated with PCOS in Hayes and Urbanek et al. (18). $P$ = significance as assessed by sample-size weighted two-strata meta-analysis, adjusted for genomic inflation.

### Subtypes in PCOS families

The RF classifier yielded the lowest overall subtype misclassification rate (13.2%) of the tested methods, according to 10-fold cross-validation of the genotyped cohort. Affected women from the family-based cohort were classified accordingly using a RF model. Seventy-three daughters of the 83 affected women from the family-based cohort had complete quantitative trait data available for subtype classification. Seventeen (23.3%) were classified as having the reproductive subtype of PCOS, and 22 (30.1%) were classified as having the metabolic subtype. Of 14 subtyped sibling pairs, only 8 were concordantly classified (57.1%); however, there was only one instance of the reproductive subtype and metabolic subtype occurring within the same nuclear family as the remaining discordant pairs each featured one indeterminate member. The proportion of affected women with one or more of the previously-identified (29) deleterious, rare variants in *DENND1A* varied by subtype. Women classified as having the reproductive subtype of PCOS were significantly more likely to carry one or more of the *DENND1A* rare variants compared to other women with PCOS (P=0.03; **Fig 11**). The distribution of affected women and *DENND1A* rare variant carriers are shown relative to the adjusted quantitative trait PCs in **Fig 12**.

**Fig 11.**
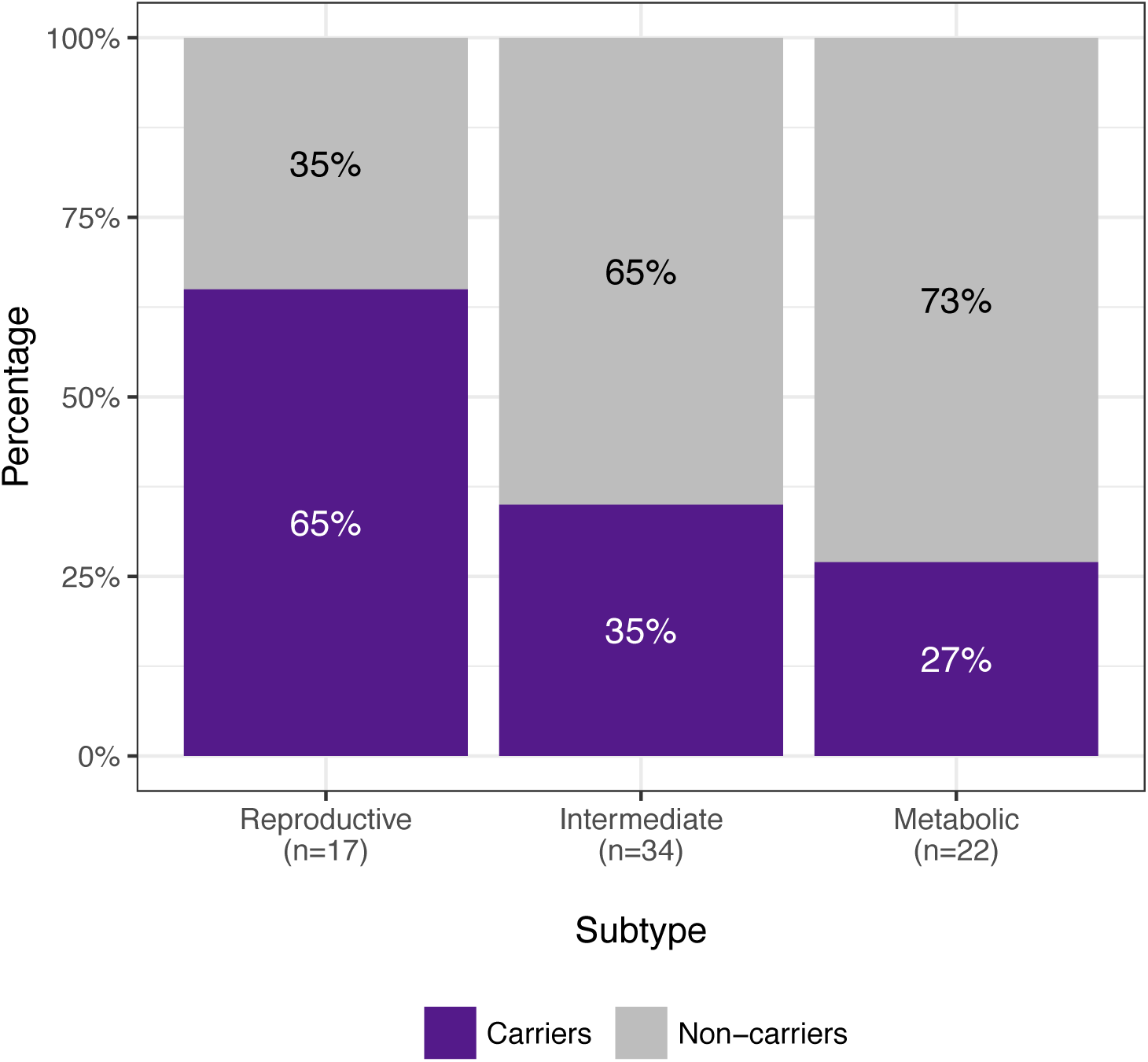
*DENND1A* rare variant carriers by subtype. The proportions of affected women with *DENND1A* rare variants in families with PCOS are shown by classified subtype. Women with the reproductive subtype were significantly more likely to carry one or more of the *DENND1A* rare variants compared to other women with PCOS (P=0.03)

**Fig 12.**
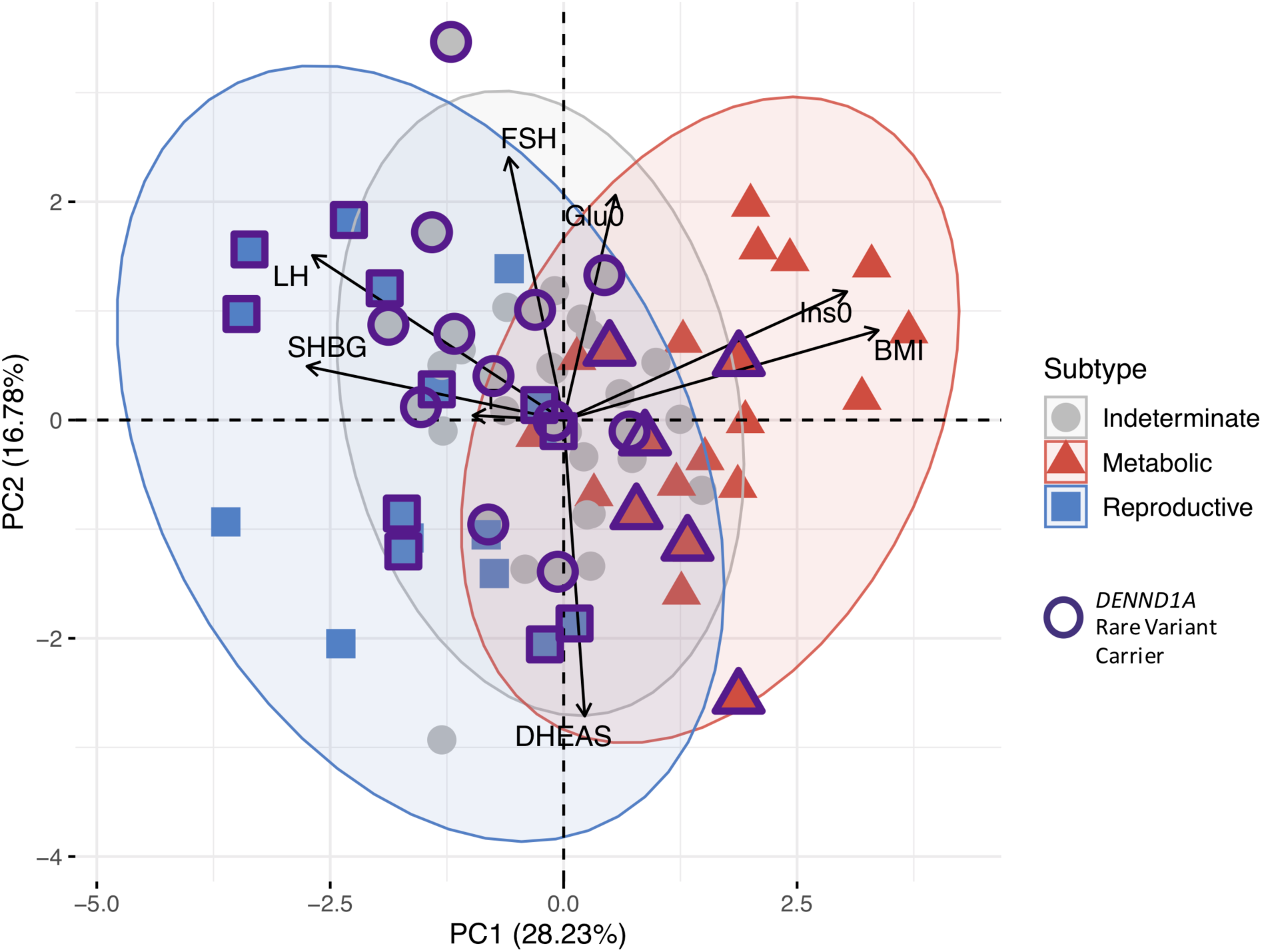
PCA of affected women in PCOS families showing *DENND1A* rare variant carriers. The distribution of affected women and *DENND1A* rare variant carriers are shown relative to their classified subtypes and their adjusted quantitative trait PCs in families affected by PCOS.

## Discussion

It is becoming increasingly clear that common, complex traits, such as T2D, are a heterogeneous collection of disease subtypes (24,45–47). There is emerging evidence that these subtypes have different genetic architectures (24,47,48). Consistent with these concepts, we identified reproductive and metabolic subtypes of PCOS by unsupervised hierarchical cluster analysis of quantitative hormonal traits and BMI and found novel loci uniquely associated with these subtypes with substantially larger effect sizes than those associated with PCOS disease status in GWAS (18–22). These findings suggest that these subtypes are both genetically distinct as well as more etiologically homogenous (9). Our findings are in contrast to the recent PCOS GWAS meta-analysis (21) that found that only one of 14 loci was uniquely associated with the NIH phenotype compared to non-NIH Rotterdam phenotypes. These latter findings suggest that the NIH and Rotterdam diagnostic criteria do not identify biologically distinct subtypes of PCOS. There have been previous efforts to subtype PCOS using unsupervised clustering (25–28), but no subsequent investigation into the biologic relevance of the resulting subtypes.

The key traits driving the subtypes were BMI, insulin, SHBG, glucose, LH, and FSH levels. The reproductive subtype was characterized by higher LH and SHBG levels with lower BMI and blood glucose and insulin levels. The metabolic subtype was characterized by higher BMI and glucose and insulin levels with relatively low SHBG and LH levels. The remaining 40% of cases had no distinguishable cluster-wide characteristics and mean trait values were between those of the reproductive and metabolic subtypes. The relative trait distributions and results of the PCAs (**Figs 3, 4, 5b**) showed the reproductive and metabolic subtypes as collections of subjects on opposite ends of a phenotypic spectrum with the remaining indeterminate subjects scattered between the two. Bootstrapping and clustering in an independent cohort revealed that the reproductive and metabolic subtypes were stable and reproducible. When the GWAS was repeated, novel susceptibility loci were associated with the reproductive and metabolic subtypes, suggesting that they had distinct genetic architecture. The indeterminate PCOS cases were associated with the reported locus at *FSHB*, but the association signal was stronger than that of our original GWAS (18), suggesting that the indeterminate group was also more genetically homogenous after the reproductive and metabolic subtypes were removed from the analysis.

Two of the loci associated with the reproductive subtype implicate novel biologic pathways in PCOS pathogenesis. The association signal on chr1 appeared downstream of and within the same TAD as the *PRDM2* gene (**Figs 8a, 9**), which is an estrogen receptor co-activator (49) that is highly expressed in the ovary (50) and pituitary gland (51). In an independent rare variant association study in PCOS families, *PRDM2* demonstrated the 5^th^ strongest gene-level association with altered hormonal levels in PCOS families (*P*=6.92×10^-3^) out of 339 genes tested (29). PRDM2 binds with ligand bound estrogen receptor alpha (ERα) to open chromatin at ERα target genes (49, 52). PRDM2 also binds with the retinoblastoma protein (53), which has been found to play an important role in follicular development in granulosa cells (54, 55).

The reproductive subtype association in the 4q22.3 locus overlapped with the *BMPR1B* gene, which transcribes a type-I AMH receptor highly expressed in granulosa cells and in GnRH neurons (16) that regulates follicular development (56). BMPR1B (bone morphogenetic protein receptor type IB), also known as ALK6 (Activin Receptor-Like Kinase 6), heterodimerizes with the TGF-β type-II receptors, including AMHR2, and binds AMH and other BMP ligands to initialize TGF-β signaling via Smads 1/5/8 (57). *BMPR1B* has been found to mediate the AMH response in ovine granulosa cells (58), and *BMPR1B*-deficient mice are infertile and suffer from a variety of functional defects in the ovary (59, 60). One of the BMPR1B ligand genes, *BMP6*, had the 3^rd^ strongest gene-level association with altered hormonal levels (*P*=4.00×10^-3^) out of 339 genes tested in our rare variant association study in PCOS families (29). Collectively, these results make *BMPR1B* a compelling candidate gene in PCOS pathogenesis. These findings also support our sequencing studies that have implicated pathogenic variants in the AMH signaling pathway in PCOS (61, 62).

The nature of the potential involvement in PCOS is less clear for the other loci associated with the reproductive subtype. The 2q37.3 locus overlapped with the promoter region of the *IQCA1* gene. Its function in humans is not well characterized, but *IQCA1* is highly expressed in the pituitary gland (51). The 5p14.2-p14.1 locus overlapped the promoter region of the *CDH10* gene (Cadherin 10). *CDH10* is almost exclusively expressed in the brain (50), and is putatively involved in synaptic adhesions, axon outgrowth and guidance (63).

The lone significant association signal with the metabolic subtype was located in an intergenic region 200-280kb downstream of the *FIGN* gene, 490-570kb upstream of *KCNH7*. *KCNH7* encodes a voltage-gated potassium channel (alias ERG3). *KCNH7* is primarily expressed in the nervous system (64), but has been found in murine islet cells (65, 66). *FIGN* encodes fidgetin, a microtubule-severing enzyme most highly expressed in the pituitary gland and ovary (50). A genetic variant in *FIGN* was found to reduce the risk of congenital heart disease in Han Chinese by modulating transmembrane folate transport (67, 68). The TAD encompassing the association signal in this locus includes *FIGN* and extends upstream to the *GRB14* gene (**Fig 9**). *GRB14* plays an important role in insulin receptor signaling (69, 70) and has been associated with T2D in GWAS (71). Given the various metabolic associations for the genes in this chromosomal region, it is plausible that causal variants in this locus could impact a combination of these genes.

Despite evidence linking neighboring genes to PCOS pathways in each of the aforementioned loci, it remains possible, of course, that other more distant genes in LD underlie the association signals. Causal variants are often up to 2 Mb away from the associated SNP, not necessarily in the closest gene (72). Fine-mapping and functional studies are needed in order to confirm the causal variants in each of these loci. In addition, the sample sizes for the subtype GWAS were small, some of the associations were based only on imputed SNPs in Stage 1, and a replication association study has not yet been performed. However, the aforementioned functional evidence for several of the loci—particularly for *PRDM2* and *BMPR1B*—support the validity of their associations. Also, the fact that one of the genes associated with the reproductive subtype, *PRDM2*, was associated with PCOS quantitative traits in our family-based analysis (29) does represent a replication of this signal by an independent analytical approach. Nevertheless, our genetic association results should be considered preliminary.

The effect sizes of the subtype alleles, particularly those associated with the reproductive subtype (Odds Ratio [OR] 3.02-5.68) (**Table 1**), were substantially greater than the effects (OR 0.70-1.51) observed for alleles associated with PCOS diagnosis in previous GWAS (18–22). In general, there is an inverse relationship between allele frequency and effect size (1) because alleles with larger phenotypic effects are subject to purifying selection and, therefore, occur less frequently in the population (73, 74). Accordingly, in contrast to the common variants (Effect Allele Frequency [EAF]>0.05) associated with PCOS in previous GWAS (18–22), the alleles associated with the subtypes were all of low frequency (EAF 0.01-0.05; **Table 1**). However, given the limited cohort sizes in this study, the subtype association testing did not have adequate power to detect associations with more modest effect sizes, such as those from our previous GWAS (18). It is also possible that the large effect sizes were somewhat inflated by the so-called “winners curse” (75, 76), but they nonetheless suggest that the subtypes were more genetically homogeneous than PCOS diagnosis in general.

In applying a subtype classifier to our family-based cohort, we found twelve affected sibling pairs in which at least one of the daughters was classified with the reproductive or metabolic subtype. Six of these sibling pairs were classified with the same subtype. There was only one discordant pairing of the reproductive subtype with the metabolic subtype. This further suggests that the reproductive and metabolic subtypes are genetically distinct in their origins. The greater prevalence of *DENND1A* rare variant carriers observed in women with the reproductive subtype in the family-based cohort implicates this gene in the pathogenesis of this subtype. *DENND1A* is known to regulate androgen biosynthesis in the ovary (77, 78); therefore, we would expect *DENND1A*-mediated PCOS to be more closely associated with the reproductive subtype of PCOS. However, we did not find an association between any *DENND1A* alleles and the reproductive subtype in the subtype GWAS, perhaps due to allelic heterogeneity or to our limited power to detect associations with more modest effect sizes.

We only studied women with PCOS as defined by the NIH diagnostic criteria. Future studies will investigate whether similar reproductive and metabolic clusters are present in non-NIH Rotterdam PCOS cases. In particular, it is possible that there will be no metabolic subtype in non-NIH Rotterdam PCOS cases since these phenotypes have minimal metabolic risk (79, 80). Indeed, in a previous effort to identify phenotypic subtypes in Rotterdam PCOS cases (28), the cluster that most closely resembled the reproductive subtype represented the largest proportion of PCOS women at 44%, of whom only 78% met the NIH criteria for PCOS, whereas the cluster that most closely resembled the metabolic subtype constituted only 12% of the cohort, but 98% met the NIH diagnostic criteria. Due to the within-cohort normalization of quantitative traits prior to clustering, our method is well-suited for identifying subsets of cases that occupy either end of a phenotypic spectrum within different populations, but it therefore may not be suitable for directly comparing subtype membership between different populations.

Our cohort included only women of European ancestry. It will be of considerable importance to investigate whether subtypes are present in women with PCOS of other ancestries. Women with PCOS of diverse races and ethnicities have similar reproductive and metabolic features (81–83). However, there are differences in the severity of the metabolic defects due to differences in the prevalence of obesity (84) and well as to racial/ethnic differences in insulin sensitivity (85, 86). Further, the susceptibility loci associated with subtypes in other ancestry groups may differ since the low frequency and large effect size of the variants associated with the reproductive subtype in our European cohort suggests these variants are of relatively recent origin, and, therefore, may be population-specific (1,87,88).

In conclusion, using an unsupervised clustering approach featuring quantitative hormonal and anthropometric data, we identified novel reproductive and metabolic subtypes of PCOS with distinct genetic architectures. The genomic loci that were significantly associated with either of these subtypes include a number of new, highly plausible PCOS candidate genes. Moreover, our results demonstrate that precise phenotypic delineation, resulting in more homogeneous subsets of affected individuals, can be more powerful and informative than increases in sample size for genetic association studies. Our findings indicate that further study into the genetic heterogeneity of PCOS is warranted and could lead to a transformation in the way PCOS is classified, studied, and treated.

## Acknowledgements

This study was supported by National Institutes of Health (NIH) Grants R01HD057223 (A.D.), P50 HD044405 (A.D.) and R01 HD085227 (A.D.). M.D. was supported by a Ruth L. Kirschstein National Research Service Award Institutional Research Training Grant, T32 DK007169. We thank the NIH Cooperative Multicenter Reproductive Medicine Network (https://www.nichd.nih.gov/research/supported/rmn) for recruiting some of the women with PCOS who participated who participated in the genomewide association study of Hayes and Urbanek et al. (18) and whose genotype data were used in this study. We also thank the following investigators for recruiting some of the control women who participated in the genomewide association study of Hayes and Urbanek et al. (18) and whose genotype data were used in this study: Dimitrios Panidis, MD, PHD (Aristotle University of Thessaloniki, Greece); Mark O. Goodarzi (Cedars-Sinai Medical Center, Los Angeles, CA); Corrine K. Welt, MD (University of Utah School of Medicine, Salt Lake City, UT; formerly of Massachusetts General Hospital, Boston, MA); Ahmed H. Kissebah (deceased, Medical College of Wisconsin, Milwaukee, WI); Ricardo Azziz, MD (State University of New York, NY; formerly of University of Alabama at Birmingham, AL); and Evanthia Diamanti-Kandarakis, MD, PhD (University of Athens Medical School, Greece).

